# Mixed fertilization strategies but not mono fertilization enhances the net primary productivity and quality of chili (*Capsicum frutescens*)

**DOI:** 10.1101/2025.08.19.671182

**Authors:** Hossain Md Dalim, Md. Golam Jilani Helal, Minhazul Kashem Chowdhury, Shoaib Rahman, Sohel Rana Mazumder, Md. Ismail Hossain, Md. Mohi Uddin Sujan Chowdury, Md. Shahariar Jaman

**Affiliations:** Ecosystem Ecology Lab, Department of Agroforestry and Environmental Science, Sher-e-Bangla Agricultural University, Dhaka, Bangladesh; Department of Horticulture, Sher-e-Bangla Agricultural University, Dhaka, Bangladesh; Department of Plant Pathology, Sher-e-Bangla Agricultural University, Dhaka, Bangladesh; Department of Agronomy, Sher-e-Bangla Agricultural University, Dhaka, Bangladesh

**Keywords:** Chili, *Leucaena leucocephala*, mixed fertilization, NPP, quality, yield

## Abstract

While the effects of different fertilization strategies on chili (*Capsicum frutescens*) cultivation have been extensively studied, the comparative assessment of mixed versus mono fertilization approaches remains still under the shadow. To address this gap, we have conducted an experiment using a completely randomized design (CRD) with five fertilization treatments along with four replications of each treatment (e.g., *Leucaena leucocephala* leaf litter and recommended dose of fertilizers (RFD)) to observed the effect of these treatments on plant productivity traits, NPP (net primary productivity), quality and overall yield of chili. Our results revealed that the mixed fertilization treatment (T₄: 100 g kg-^1^ L*. leucocephala* + ½ RFD) produced the highest yield (15.50 t ha⁻¹), a 63.9% increase over the control. We also found that plant height (73.75 cm), number of leaves (1000.25 plant^-1^), number of fruits (93.00 plant^-1^) were higher at 60 DAT (days after transplanting) however, number of flowering (44.50 plant^-1^) at 45 DAT. Similarly, Above- and belowground NPP (26.47 g plant^-1^), vitamin C (124.65 mg 100 g⁻¹ ), capsaicin content (1.22 %) and SPAD chlorophyll index (48.61) were higher in T_4_ treatment compared to T_0_ treatment. Furthermore, regression results showed that ANPP and yield significantly increased with increase the number of leaves (ANPP: *R²* = 0.51, *p* = 0.02; Yield: *R²* = 0.34, *p* = 0.002) and number of flowering (ANPP: *R²* = 0.41, *p* = 0.014; Yield: *R²* = 0.42, *p* = 0.037) as well. Our results suggest that mixed fertilization enhances both productivity and quality traits in chili. Therefore, promotion of integrated (organic and inorganic together) nutrient management practices is recommended to improve yield and quality of chili.

## 1. Introduction

Chili (*Capsicum frutescens*) is widely used spice belongs to the *Capsicum* genus of the *Solanaceae* family [1] and now extensively cultivated around the world [2]. It is indigenous to the America and West Indies but began its global spread to tropical regions after those areas were explored [3]. Bangladesh is not an exception where chili cultivation encompasses a significant portion of the country’s agricultural landscape. For instances, during the 2022–2023 chili was cultivated on approximately 188,888.78 ha of land in Bangladesh and producing about 507,200.88 metric tons of yield [4]. As global consumption of chili increases, the demand for improved fruit quality and overall productivity has increased [5] which largely depended on intensive agricultural practices particularly the application of inorganic fertilizers [6,7]. Although inorganic fertilization boost up vegetative and reproductive development of chili, alternatively, it negatively affect the soil health for long term [8]. Thus, application of organic fertilizers along with minimum use of inorganic fertilizes would be a better alternative choice for chili productivity which have not yet sufficiently explored and crucially need to investigate.

Traditionally, mono fertilization either through chemical (inorganic) or organic means has been a predominant method used by farmers. Inorganic fertilizers are known for their immediate nutrient availability and are widely employed to boost crop productivity [9,10]. However, long-term use of chemical fertilizers alone can lead to nutrient imbalances, soil acidification and reduced microbial diversity, potentially compromising soil health and crop quality [11,12]. On the other hand, organic fertilizers such as compost, manure and green residues enhance soil structure, increase organic matter content and support microbial activity but their nutrient release is often slower and less predictable [13,14]. Hence, relying solely on mono fertilization may not effectively meet the nutritional demands of high-yielding chili varieties or address the sustainability concerns of modern agriculture [15]. Mixed fertilization strategies which integrate both organic and inorganic nutrient sources have been increasingly recognized as a holistic and sustainable approach to soil fertility and crop management. These integrated nutrient strategies aim to exploit the rapid nutrient availability of inorganic fertilizers, while simultaneously harnessing the long-term soil enhancement provided by organic amendments such as green manures and composts [16]. Specifically, the incorporation of *Leucaena leucocephala* leaf litter alongwith inorganic fertilizers has shown potential to enhance crop productivity and soil health by synchronizing nutrient release with crop demand, reducing reliance on synthetic inputs and promoting ecological sustainability [17,18]. Such practices are believed to improve soil physicochemical properties and support a more robust soil microbiome [19]. Consequently, it is hypothesized that mixed fertilization not only contributes to increased net primary productivity (NPP) but also improves the nutritional composition and market quality of chili (*Capsicum frutescens*) [20,21]. Integrating organic and inorganic nutrient sources can significantly enhance plant yield contributing parameters. For instance, the combined application of urea at 90 kg N ha⁻¹ with poultry manure at 30 kg N ha⁻¹ resulted in marked improvements in vegetative traits, including a 26% increase in plant height e.g., 30% expansion in leaf area and a 32% rise in the number of leaves per plant. Likewise, fruit weight, postharvest quality and nitrogen uptake efficiency improved by 36%, 39% and 50% respectively [22].

In mixed fertilization treatments the concurrent benefits of mineral nutrient availability and organic matter enrichment create a favorable environment for improved physiological performance and metabolic processes. Malik et al. [23] demonstrated that combined use of urea and farmyard manure significantly increased both the dry matter accumulation and vitamin C content in peppers. Similarly, integrating vermicompost with recommended doses of NPK fertilizers has been shown to significantly enhance the quality of *Capsicum* by increasing levels of carotenoids, capsaicinoids, vitamin A, vitamin C and capsaicin content [24,25]. The combined application of 75% of the recommended dose of fertilizer (RDF) and organic manure notably improved chlorophyll content in chili leaves throughout the crop’s growth stages [26]. The application of farmyard manure blended with inorganic fertilizer significantly increased the total fresh fruit yield of chili compared to unfertilized plots [27]. Generally, use of synthetic fertilizer such as N, P, K increase above ground biomass productivity through instant soil fertility enhancement and ion exchange capacity, interestingly, however, when combined with organic amendment it drives plant below ground biomass productivity as well. Therefore, these together approaches have been proved to provide overall productivity benefits for plants [28]. Vegetative attributes like leaf count and flowering along with reproductive factors such as fruit number and size collectively influence the above ground net primary productivity (ANPP) and final yield. Yield is the cumulative expression of various growth and physiological processes many of which are influenced by nutrient-driven changes in ANPP. For example, greater leaf area and branching improve light interception and photosynthetic surface thereby raising ANPP and ultimately leading to higher fruit yield [29]. Furthermore, efficient nutrient use improves internal plant nutrient partitioning and carbohydrate metabolism both of which are linked with increased fruit biomass and improved harvest index [30]. Several studies have explored the impact of integrated nutrient management on various crops, showing that a balanced input of organic and inorganic fertilizers can lead to higher yields, better nutrient uptake and improved crop quality [31–33]. However, comprehensive investigations specifically focusing on *Capsicum frutescence* remain limited. Moreover, the effects of mixed fertilization on critical parameters such as fruit size, capsaicin content, vitamin C concentration and overall biomass production in chili have not been sufficiently addressed. To do so, we conducted this experiment to analyze the effect of mixed and mono fertilization on potential yield contributing characters of chili and to find out the effect of mixed and mono fertilizer on NPP and quality of chili. Moreover, the research would provide insight on the relationship between potential yield contributing character with ANPP and yield of chili. Therefore, our specific objectives are (i) To analyze the effect of mixed and mono fertilization on potential yield contributing characters of chili; (ii) To find out the effect of mixed and mono fertilizer on NPP and quality of chili and (iii) To evaluate the relationships between potential yield contributing character with ANPP and yield of chili.

## 2. Materials and methods

### 2.1. Experimental site and plant material

The experiment was conducted from July to December 2023 at the research farm of the Department of Agroforestry and Environmental Science, Sher-e-Bangla Agricultural University, Dhaka-1207, Bangladesh. The site is geographically positioned at 23.77°N latitude and 90.35°E longitude, with an elevation of approximately 8.6 meters above sea level. The region experiences a subtropical monsoon climate with significant rainfall during the monsoon and moderate temperatures year-round. During the experimental period the average monthly temperature ranged from 23.8 °C to 31.4 °C while the average relative humidity fluctuated between 67% and 87%. The total rainfall recorded was approximately 1100 mm primarily concentrated in July and August (Fig 1). Sunshine hours varied between 4.3 and 7.2 hours per day [34].

**Fig 1.**
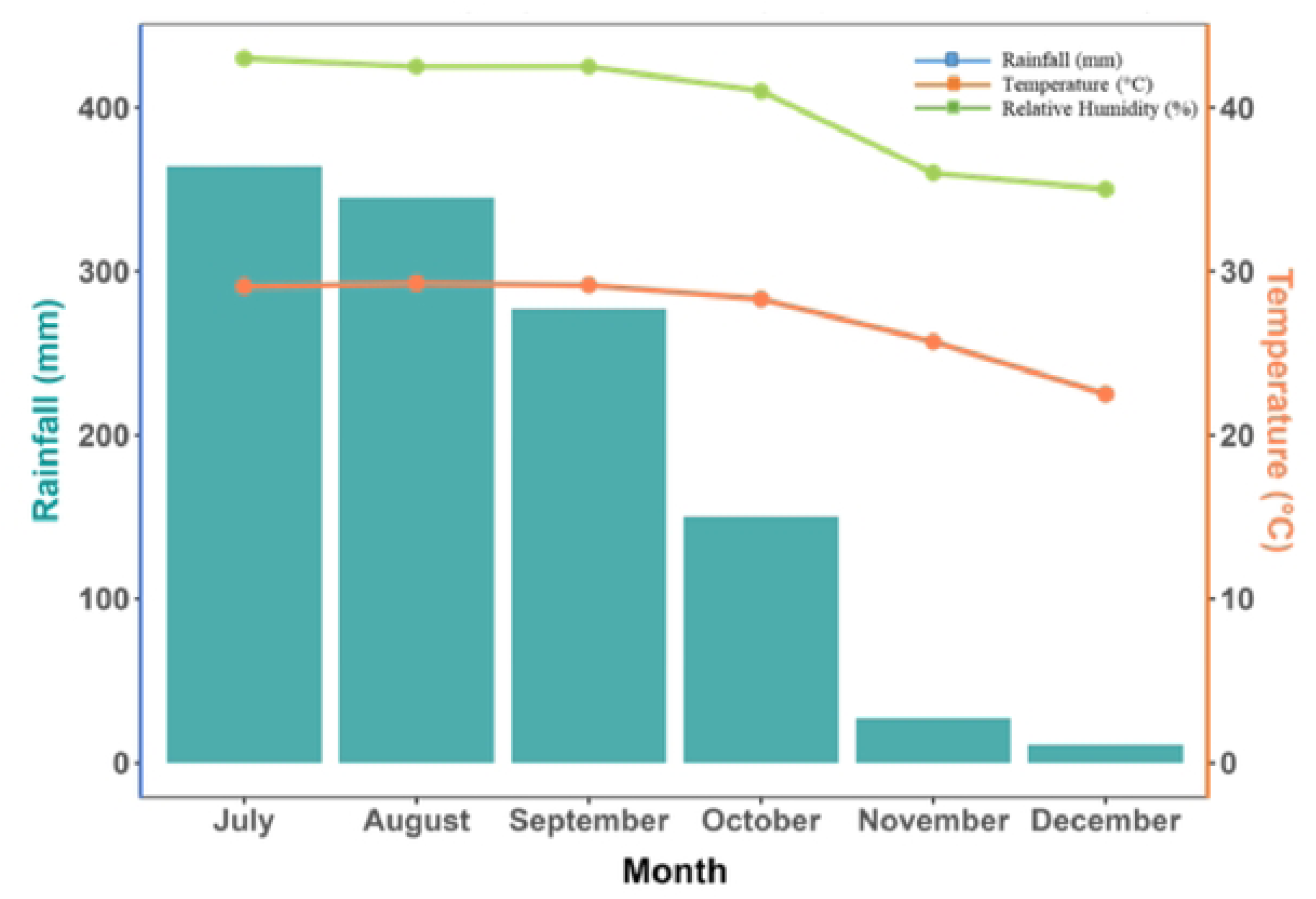
Meteorological data (rainfall (mm), temperature (°C) and relative humidity (%)) during the experiment period (July–December 2023) derived from BMD (Bangladesh Meteorological Department) Sher-e-Bangla Nagar, Dhaka.

The experimental soil was classified as silty loam with good drainage and moderate fertility. Composite soil samples were collected before the experiment. The plant material used was chili (*Capsicum frutescens*), variety BARI-3, obtained from the Bangladesh Agricultural Research Institute (BARI). This variety is known for its adaptability, pest resistance, and high yield potential. Seedlings were raised in a nursery and transplanted into pots 20 days after sowing with an average seedling height of 15 cm at transplanting.

### 2.2. Pot preparation and experimental design

The experiment was laid out in a completely randomized design (CRD) comprising five fertilization treatments each replicated four times for a total of 20 pots (Fig 2). Plastic pots measuring 30 cm in height and 28 cm in diameter (approximately 18–20 liters capacity) were used. Each pot was thoroughly cleaned, perforated at the base for drainage, and a gravel layer was added to prevent waterlogging. Topsoil from the upper 0–20 cm layer was collected, air-dried, sieved (2 mm mesh), and homogenized to ensure uniform texture and fertility. Each pot was filled with 8 kg of prepared soil.

**Fig 2.**
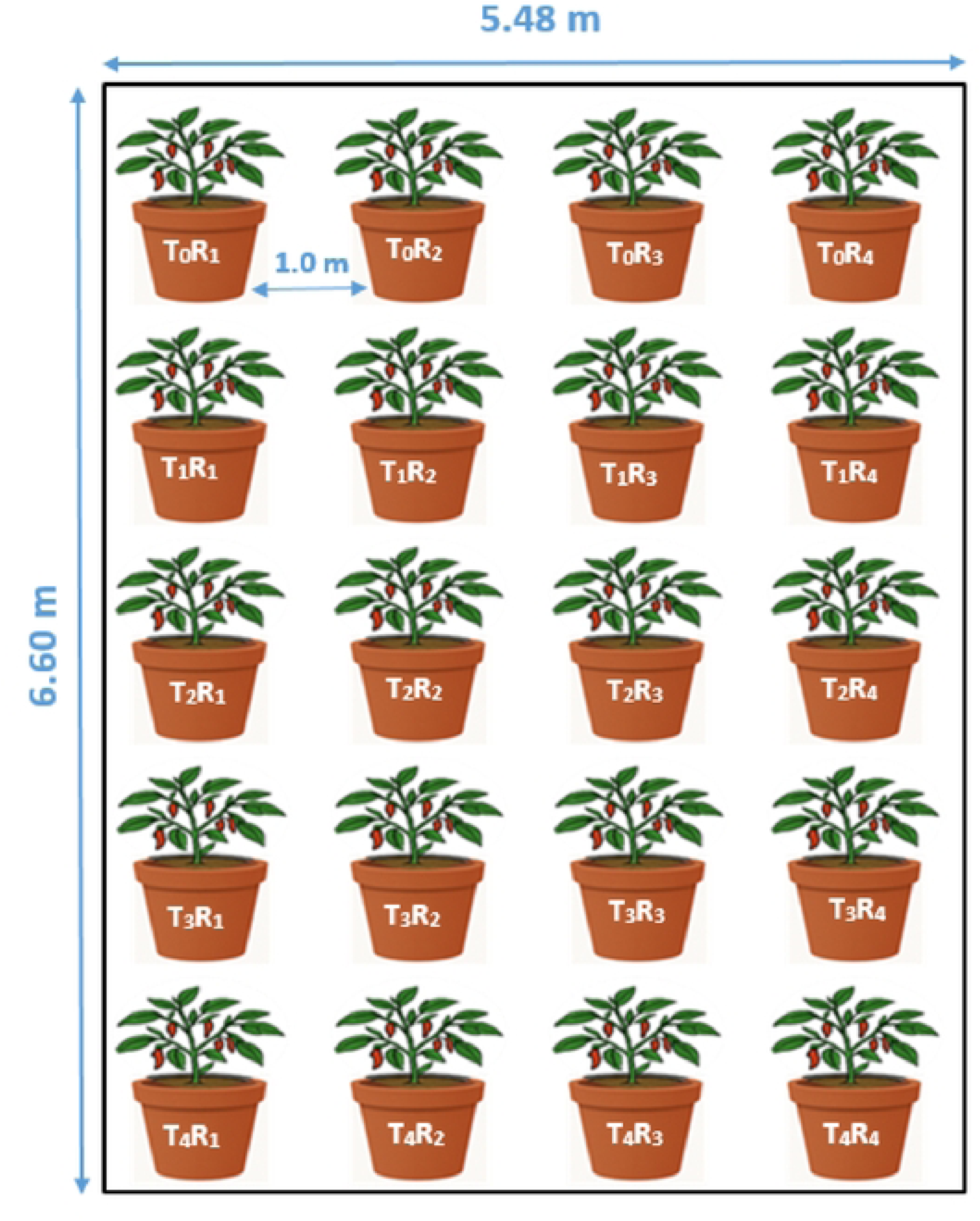
Field lay out of the experimental plot.

Inorganic fertilizers (urea, TSP, MOP, gypsum, boric acid and zinc sulfate) and organic material (oven-dried *Leucaena leucocephala* leaf litter) were incorporated into the soil one week before transplanting according to treatment specifications. After incorporation the soil was stabilized and one healthy 20-day-old chili seedling (BARI-3 variety) was transplanted into each pot. Pots were arranged randomly with 1.0 m spacing to ensure uniform sunlight exposure and airflow. All pots were labeled clearly and crop management practices were applied uniformly across treatments throughout the experiment. Standard agronomic practices were uniformly applied across all pots.

Regular watering was performed to maintain field capacity and manual weeding was done as needed. Pest and disease control measures were implemented using recommended safe practices to ensure uniform crop health throughout the growing season.

Before treatment application, composite samples were analyzed for pH and key nutrient contents. Parallel nutrient analysis was carried out for the *Leucaena leucocephala* leaf litter intended for organic amendments (Table 1).

**Table 1.** Nutrient composition of the experimental soil before treatment application and nutrient composition of the decomposed *Leucaena leucocephala* leaf litter.

| Nutrient parameter | Unit | *Leucaena leucocephala* leaf litter | Initial soil before experiment |
| --- | --- | --- | --- |
| pH | — | 6.9 | 6.5 |
| Total Nitrogen (N) | % | 1.19 | 0.08 |
| Phosphorus (P) | mg/kg soil | 0.16 | 92.72 |
| Potassium (K) | mEq/100g | 0.04 | 0.66 |
| Sulfur (S) | mg/kg soil | 0.15 | 46.14 |
| Magnesium (Mg) | % (leaf) / mEq/100g (soil) | 0.48 | 9.48 |
| Boron (B) | mg/kg soil | 2.43 | 0.78 |

#### 2.2.1. Fertilizer application

Fertilizer treatments combined with organic and/or inorganic nutrient sources in different proportions are shown in Table 2. The inorganic fertilizers e.g., urea, triple superphosphate (TSP), muriate of potash (MOP), gypsum, boric acid and zinc sulfate-were applied either at the full recommended dose (RFD) or at half the RFD, calculated according to national agronomic guidelines and converted to per-pot quantities.

**Table 2.** Rates of different fertilizer doses varied by treatments.

| **Treatment** | **Organic fertilizer** | **Inorganic fertilizer\*** |
| --- | --- | --- |
| T<sub>0</sub> | None | RFD |
| T<sub>1</sub> | *L. leucocephala* leaf litter- 12.4 g kg<sup>-1</sup> soil (93 g pot<sup>-1</sup>) | None |
| T<sub>2</sub> | *L. leucocephala* leaf litter- 24.8 g kg<sup>-1</sup> soil (186 g pot<sup>-1</sup>) | None |
| T<sub>3</sub> | *L. leucocephala* leaf litter- 12.4 g kg<sup>-1</sup> soil (93 g pot<sup>-1</sup>) | ½ RFD |
| T<sub>4</sub> | *L. leucocephala* leaf litter- 24.8 g kg<sup>-1</sup> soil (186 g pot<sup>-1</sup>) | ½ RFD |

\***RFD**- Urea (1.0 g), TSP (1.10 g), MOP (0.90 g), Gypsum (0.60 g), Boric acid (0.25 g) and zinc sulfate (0.25 g). ½ **RFD**- Urea (0.50 g), TSP (0.55 g), MOP (0.45 g), Gypsum (0.30 g), Boric acid (0.13 g), Gypsum (0.13 g) and zinc sulfate (0.13 g) [Fertilizer application rates were calculated and converted to g pot<sup>-1</sup> according to RFD regulation of t ha<sup>-1</sup>] [35–37]

#### 2.2.2. Application of litterbag technique

To investigate the decomposition dynamics and nutrient release patterns of L. *leucocephala* leaf litter the litterbag technique was employed a widely accepted method for studying organic matter breakdown in agroecological [38,39]. Nylon mesh bags (2 mm) containing 10 g of air-dried litter were buried 5 cm deep in relevant treatment pots to allow microbial access while retaining material integrity [40]. Litterbags were retrieved at 15-day intervals over 60 days. After collection the litter was gently washed to remove adhering soil particles then oven-dried at 65 °C until a constant weight was reached and subsequently weighed to determine mass loss as an indicator of decomposition and nutrient mineralization[41]. Chemical analysis of the decomposed litter and surrounding soil revealed significant nutrient enrichment. The results showed a measurable release of key nutrients (Table 2). These findings highlight the role of *L. leucocephala* litter in enhancing soil fertility and supporting plant growth through sustained nutrient release.

### 2.3. Data collection

In this study various growth, yield, productivity and quality parameters of chili (*Capsicum frutescens*, var. BARI-3) were measured using standardized procedures. Details for each parameter are provided below:

Plant growth was monitored through regular measurements of plant height and leaf count at 15, 30, 45, and 60 days after transplanting. Plant height was measured from the soil surface to the apical tip of the main stem using a standard ruler. Fully developed leaves were counted manually for each plant, excluding senescent and immature leaves. Flowering and fruiting were recorded at 30 and 45 days after transplanting, with a final fruit count taken at 60 days to monitor full crop development. All fully opened flowers and mature fruits were counted manually per plant and averaged across replications. Aboveground Net Primary Productivity (ANPP) was assessed at maturity by harvesting all shoot biomass (leaves and stems), which was oven-dried at 60 °C for 48 hours to constant weight and expressed in g plant^-1^[42]. BNPP was determined by carefully uprooting plants and placing them in plastic bags, stored at 4 °C until processing. Sampling was done up to a 10 cm soil depth to recover most root biomass. Roots were separated from soil by gently washing under running water through a 2 mm mesh sieve to collect both fine and coarse roots. Cleaned roots were oven-dried at 60 °C for 48 hours or until a constant weight was reached. After drying, samples were cooled in a desiccator and handled with clean, dry forceps. The dried roots were then transferred into labeled brown paper envelopes. Final root biomass was measured using a precision digital balance and recorded as dry root biomass in g plant⁻¹.[42]. Fruit yield per plant was recorded at final harvest using a precision digital balance. The yield per hectare (t ha^-1^) was extrapolated from per plant yield using the formula [43]:

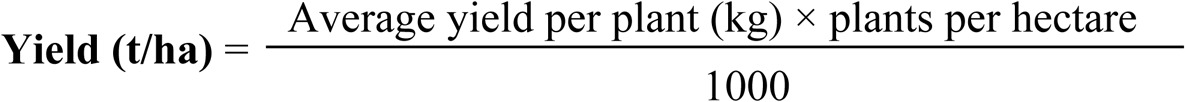

Vitamin C content in chili fruits was quantified using the 2, 6-dichloroindophenol titration method [44]. Samples were heat-treated at 70 °C, dried to constant weight, powdered, and then titrated to determine ascorbic acid concentration, expressed as mg 100 g^-1^ fresh weight. Capsaicin content was determined colorimetrically using a (Thermo, GENESYS 10 UV) spectrophotometer. Absorbance was measured at 286 nm and compared against a standard capsaicin calibration curve (0-0.10 mg mL^-1^) prepared in a methanol: ethanol: water (6:2:2 v/v) solution [45]. The results were expressed as mg capsaicin g^-1^ dry sample. Chlorophyll content was estimated using a SPAD meter (SPAD-502; Konica Minolta Sensing, Inc., Osaka, Japan) was used to measure chlorophyll content in leaf of chili. For each plant, three leaves were randomly selected and SPAD readings taken at their midpoints were averaged.

### 2.4. Data analysis

All collected data were subjected to rigorous statistical analysis to evaluate the effects of mixed and mono fertilization on the growth, productivity, and quality traits of *Capsicum frutescence*. The analysis was performed using R software (version 4.3.1; R Core Team, 2023)[46] and figures were generated using the ggplot2, tibble, and plyr packages [47]. Additional R packages, such as ggpubr [48], ggpmisc [49], gridExtra [50] and nlme [51], were also utilized for graph plotting. Prior to hypothesis testing data were assessed for normality and homogeneity of variance using the Shapiro-Wilk test [52] and Levene’s test [53] respectively to ensure compliance with the assumptions of parametric analysis. Variables that met these assumptions were subjected to one-way analysis of variance (ANOVA) to determine the significance of treatment effects across all measured parameters including morphological (plant height, leaf number), reproductive (flower and fruit number), biomass productivity (ANPP, BNPP), yield and fruit quality traits (vitamin C content, capsaicin concentration and chlorophyll index). Where ANOVA indicated significant differences, treatment differences and average values were partitioned with Tukey’s Honestly Significant Difference (HSD) test where the differences were predicted at *p* < 0.05 significant level [54]. Next, linear fit (*R^2^*) regressions analysis were performed with the ‘lm’ function to see the bivariate relation (e.g., number of leaves per plant, number of flowering per plant, with ANPP and chili yield) and significance were indicated at *p* < 0.05, *p* < 0.01 and *p* < 0.001 level. We used F-statistics to determine whether the variance between two standard variables is similar.

## 3. Results

### 3.1 Mixed fertilization significantly enhanced growth dynamics

Plant height and leaf number were progressively influenced by fertilization regimes across all observation periods. At 15 DAT differences among treatments were minimal (Figs 3a and 4a). However, at 30 DAT, plants under mixed fertilization (T₄) had begun to show enhanced growth reaching (mean ± se: 56.75 ± 6.59 cm) in height (Fig 3b) and (652.00 ± 21.67) leaves plant^-1^ (Fig 4b), compared to (41.00 ± 4.20 cm; *p* = 0.04) and (468.50 ± 36.27; *p* < 0.001) leaves in the control (T₀). These differences became more pronounced at 45 DAT with T₄ achieving (66.75 ± 7.60 cm; *p* = 0.03) height (Fig 3c) and (849.25 ± 25.89; *p* < 0.001) leaves (Fig 4c) maintaining a clear advantage. At 60 DAT, the growth response peaked in T₄ (73.75 ± 7.87 cm; *p* = 0.02) (Fig 3d) and (1000.25 ± 19.08; *p* < 0.001) leaves (Fig 4d), while other treatments remained significantly lower.

**Fig 3.**
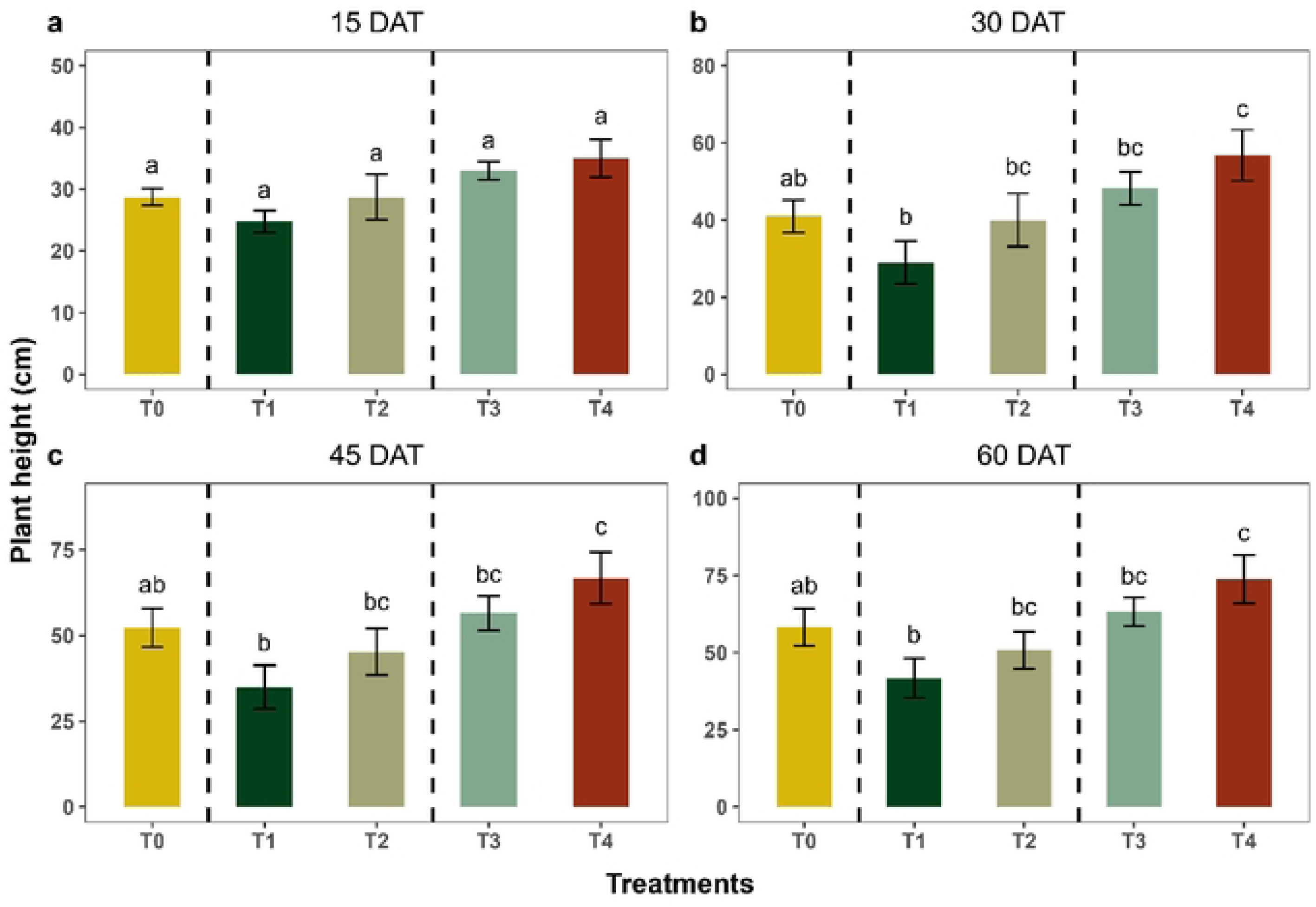
Effect of different fertilization treatments on plant height of *Capsicum frutescence* at various DAT.

**Fig 4.**
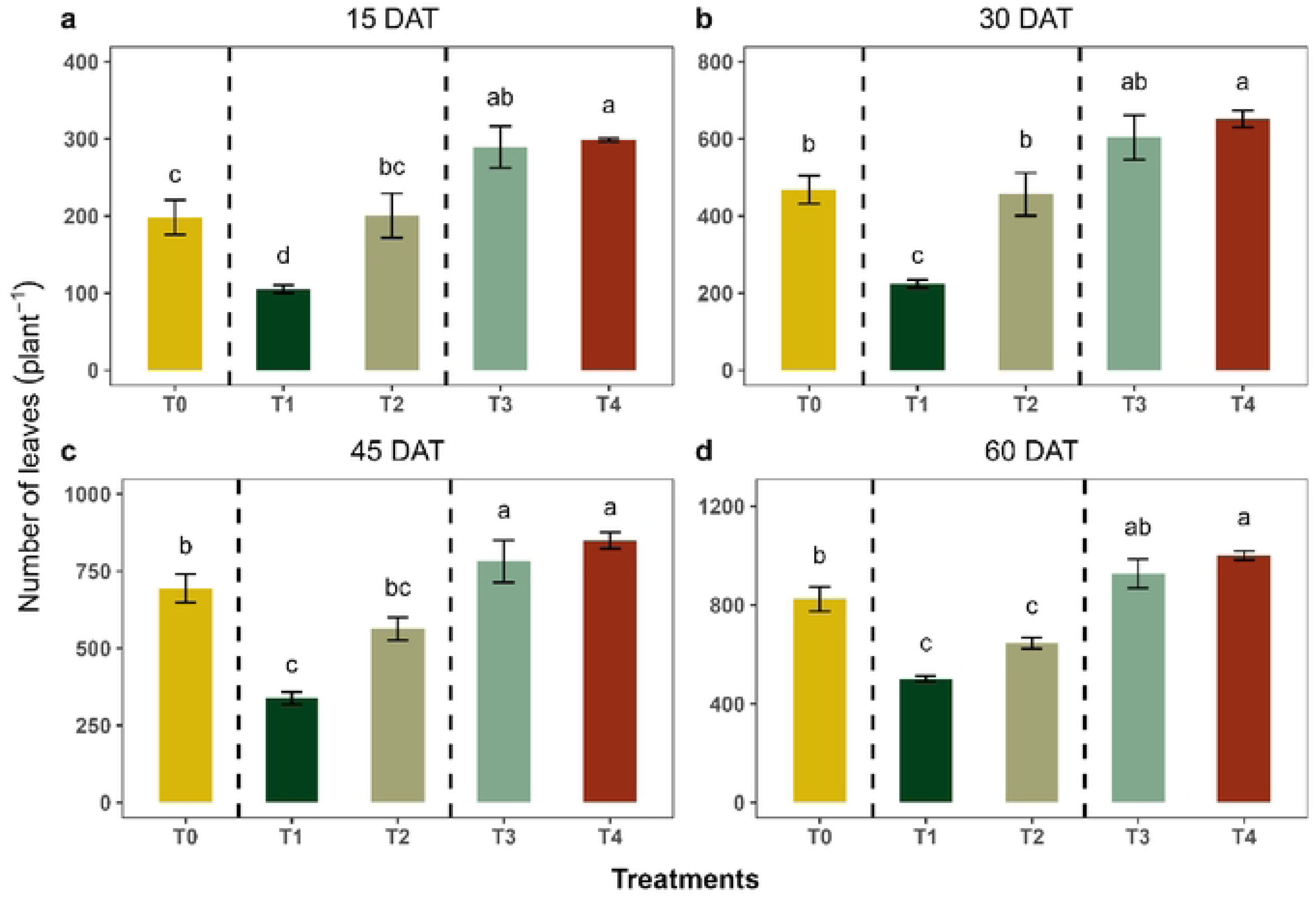
Effect of different fertilization treatments on the number of leaves per *Capsicum frutescence* plant at various DAT.

### 3.2 Enhanced reproductive traits under mixed fertilization

Reproductive traits including flower and fruit number varied significantly among treatments and across growth stages. At 30 DAT, flower production was highest in T₄ (22.50 ± 4.11; *p* = 0.01) followed by T₃ (22.25 ± 1.65) (Fig 5a), while number of flowerings at 30 DAT showed no significant differences among T_0_, T_1_ and T_2_ treatments respectively. Flower number increased notably at 45 DAT in T₄ (44.50 ± 9.47; *p* = 0.03) and T₃ (38.50 ± 1.55) treatments respectively, whereas T_0_, T_1_ and T_2_ showed no significant variation (*p* > 0.05) (Fig 5b). When considering fruit, we found that fruit formation began at 30 DAT (Fig 6a) but significant results were observed at 45 DAT, in which T₄ produced (43.0 ± 2.42; *p* < 0.001) fruits plant^-1^ compared to control (16.5 ± 2.33) (Fig 6b). At 60 DAT, T₄ treatment showed significantly higher number of fruits plant^-1^ (93.00 ± 5.49), followed by T₃ (59.25 ± 5.53) and T₂ (43.00 ± 4.10) respectively (Fig 6c) while T₀ exhibited (47.25 ± 7.19) fruits plant^-1^, which was significantly less than the T₄ treatment (*p* < 0.001).

**Fig 5.**
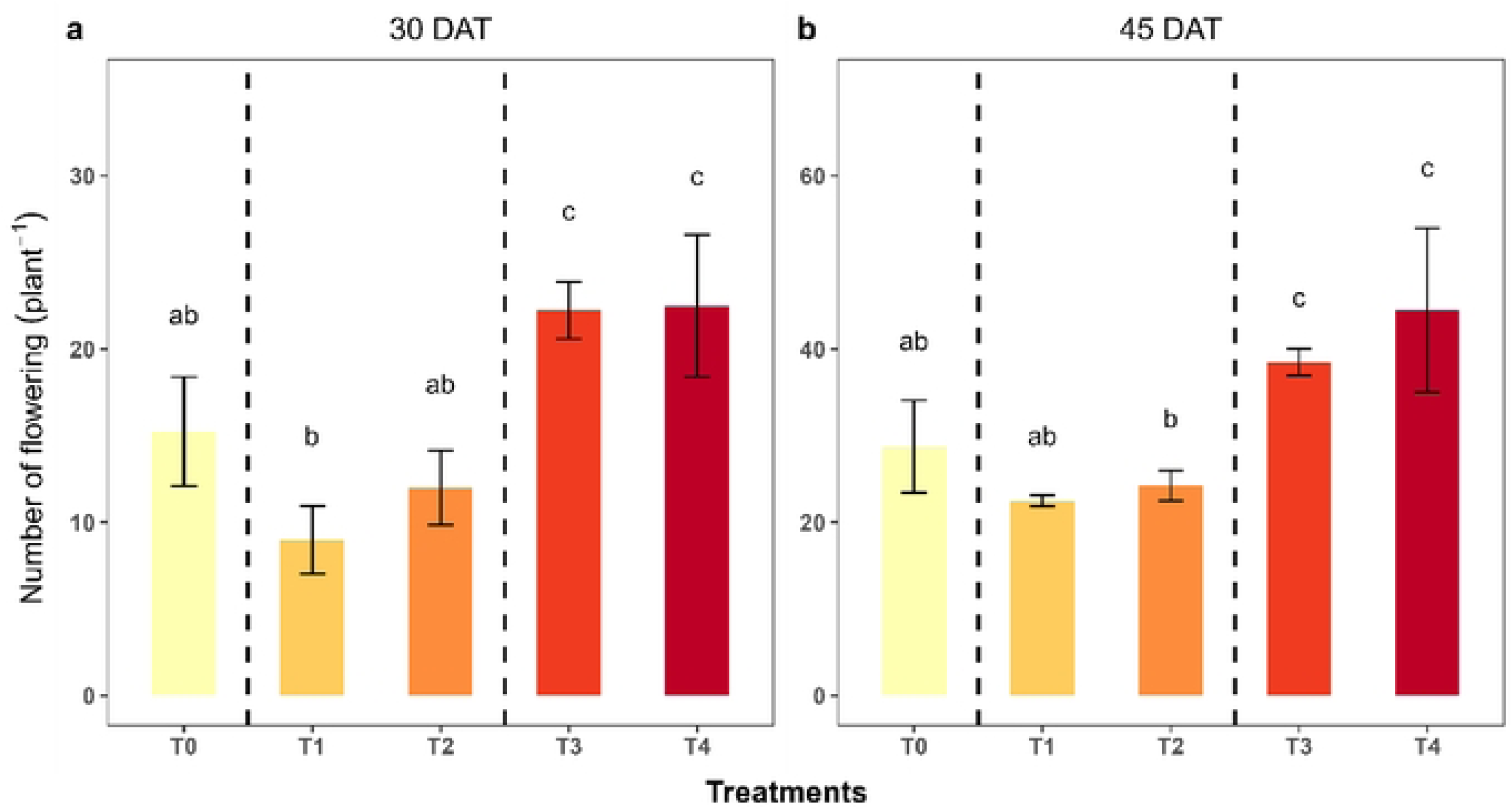
Number of flowers plant^-1^ under different fertilization treatments at 30 and 45 DAT.

**Fig 6.**
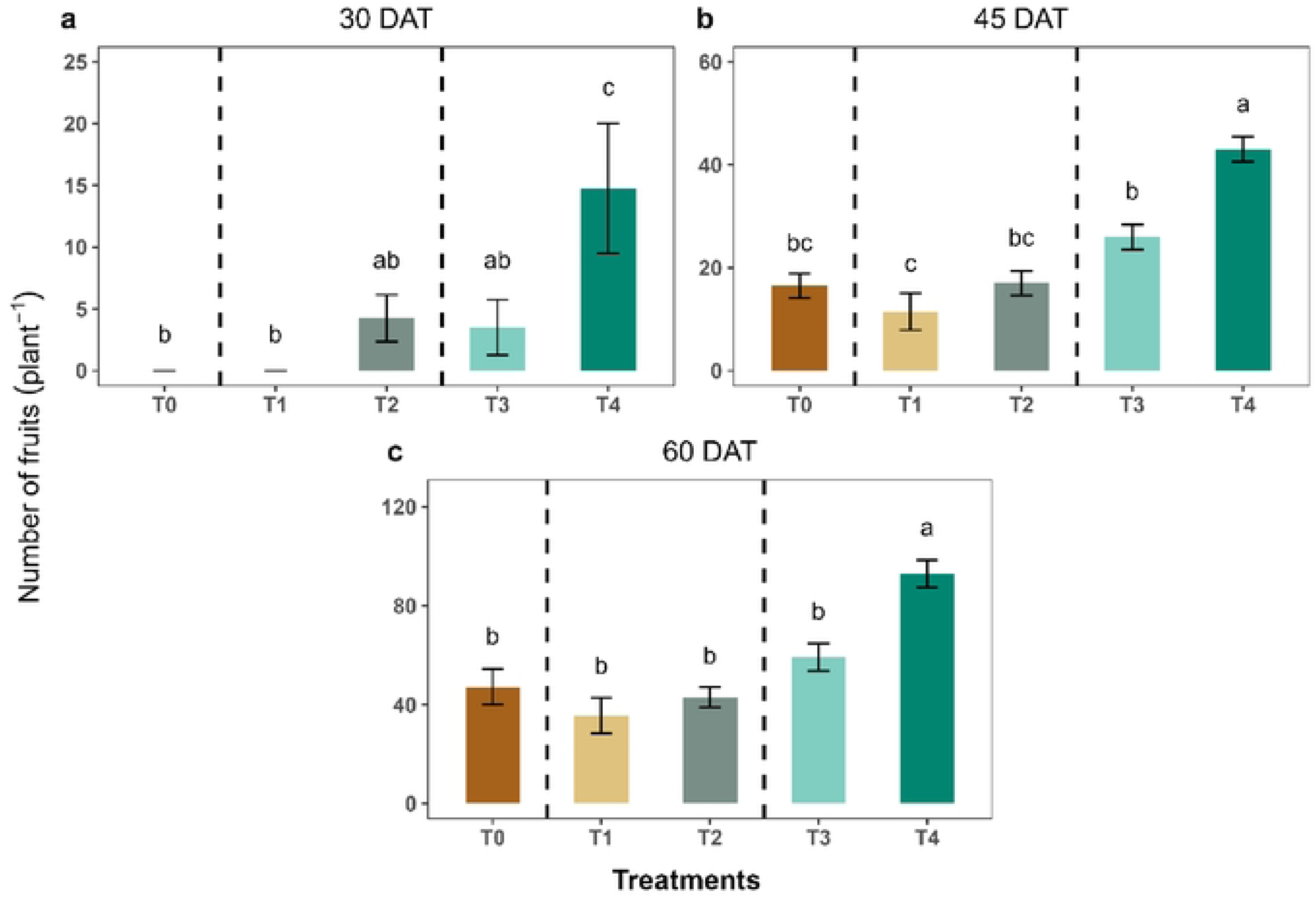
Number of fruits plant^-1^ under different fertilization treatments at 30, 45 and 60 DAT.

### 3.3 Net primary productivity maximized under mixed inputs

Our results revealed a significant effect on NPP varied by different treatment. ANPP in mixed fertilizer (T_3_ + T₄) treatments showed (21.82 ± 2.20 g plant^-1^; *p* < 0.001), representing 26.67% and 116.39% increase compared to inorganic (T_0_) and organic (T_1_ + T_2_) fertilizer respectively (Fig 7a). Root biomass (BNPP) enhanced by mixed fertilization reaching (3.93 ± 0.39 g plant^-1^) under (T_3_ + T₄) treatments, which was significantly enhanced over mono fertilization (*p* < 0.001) (Fig 7b). Total NPP (ANPP + BNPP) was found higher in (T_3_ + T_4_) treatments (26.47 ± 3.16 g plant^-1^) followed by (T_1_ + T_2_) treatments (13.84 ± 1.37 g plant^-1^) and control (20.48 ± 2.47 g plant^-1^; *p* = 0.01682) respectively (Fig 7c).

**Fig 7.**
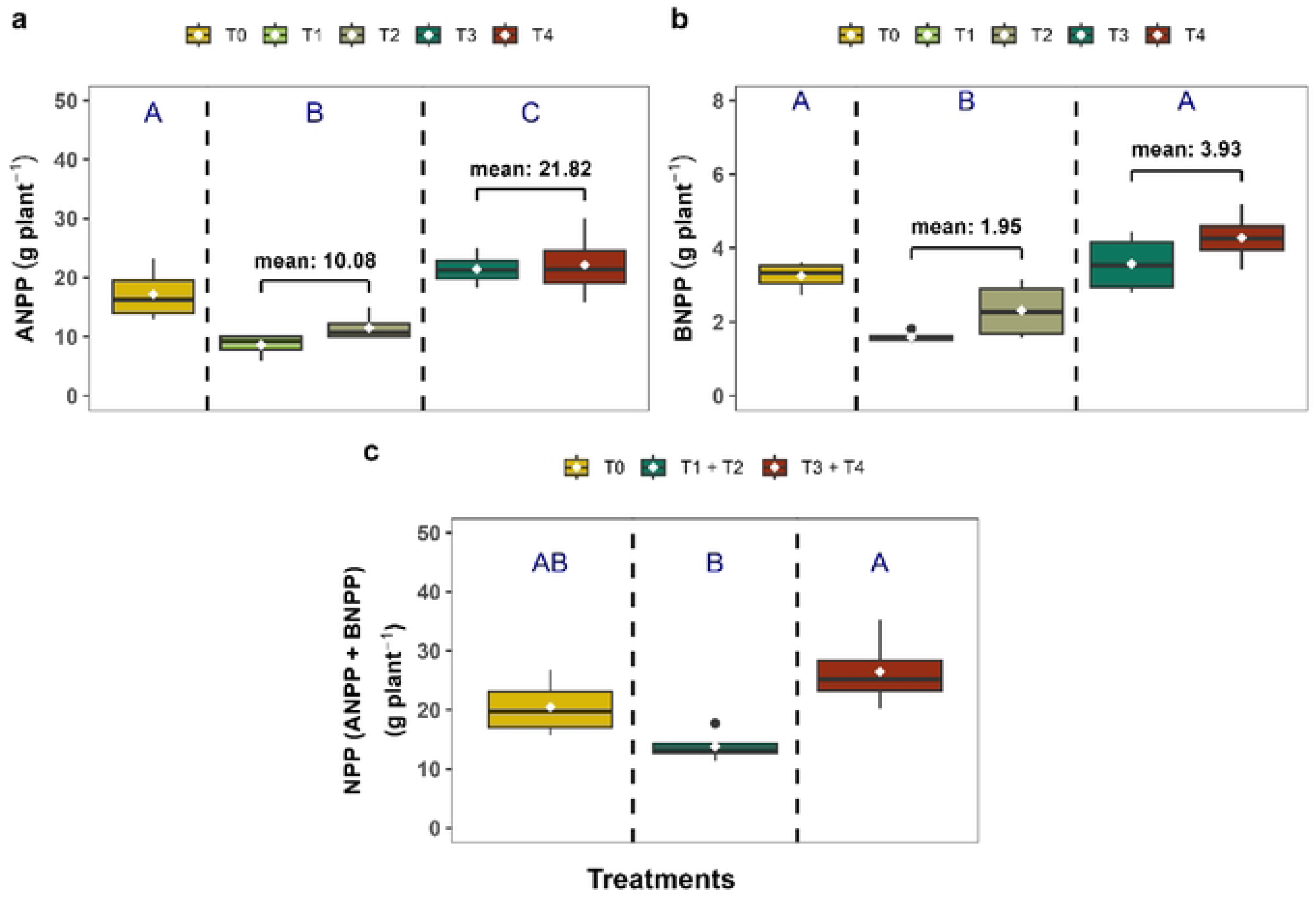
Net primary productivity (ANPP + BNPP) of *Capsicum frutescence* under different fertilization treatments.

### 3.4 Mixed fertilization significantly boosted fruit yield

Yield was notably improved under the mixed fertilization treatments (T₃ + T₄). Mixed fertilizer produced (13.01 ± 1.24 t ha^-1^; *p* = 0.003) fruits plant^-1^, which was 37.48% higher than the inorganic fertilizer (T_0_) and 60.29% above organic fertilizer (T_2_ + T_4_) treatments (Fig 8). The application of organic and inorganic fertilizers exhibited clear significant differences with mixed fertilizer application and has no significant differences between them. T₄ treatment produced (15.50 ± 0.54 t ha^-1^) fruit**s** plant^-1^, which was 63.90% higher than the control, as well as 106.21% above T_1_ and 78.02% over T_2_ treatments respectively. Yield was also showed higher in T₃ treatment (10.51± 1.92 t ha^-1^), though slightly less than T₄ (Fig 8).

**Fig 8.**
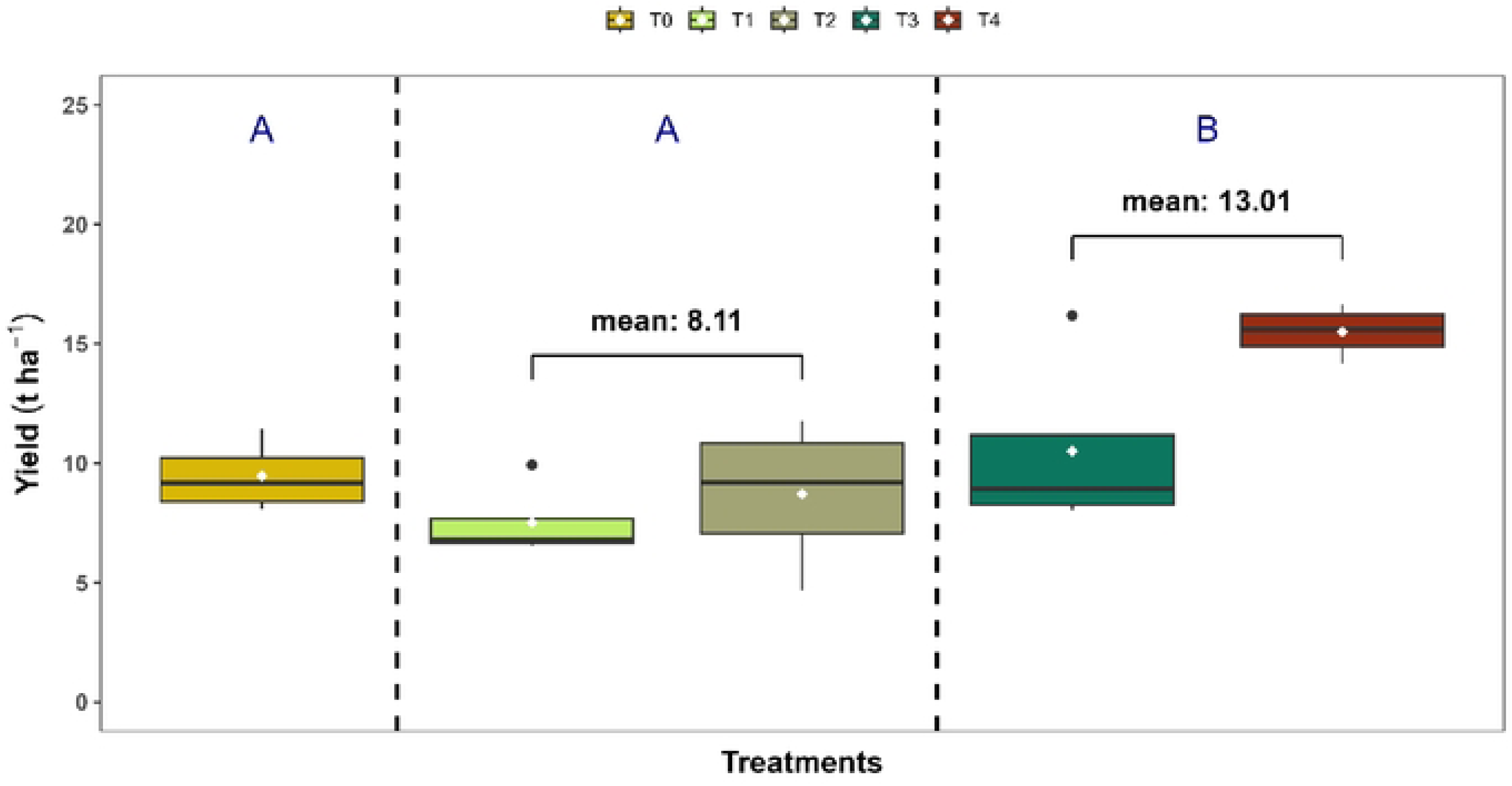
Fruit yield (t ha^-1^) of *Capsicum frutescence* under different fertilization treatments.

### 3.5 Quality attributes significantly elevated with integrated nutrients

The influence of fertilization on quality parameters such as vitamin C, capsaicin content and chlorophyll index showed in figure 9. Fruit under treatment T₄ recorded as highest vitamin C content (124.65 ± 1.39 mg 100 g^-1^) outperforming all other treatments significantly (Fig 9a). Capsaicin content was higher in T₄ treatment (1.22 ± 0.01 %) which was found to be increased 1.07 ± 0.01 % over the control (Fig 9b). Moreover, leaf SPAD chlorophyll index was highest in T₄ (48.61 ± 0.14) indicating better nitrogen assimilation and photosynthetic efficiency (Fig 9c).

**Fig 9.**
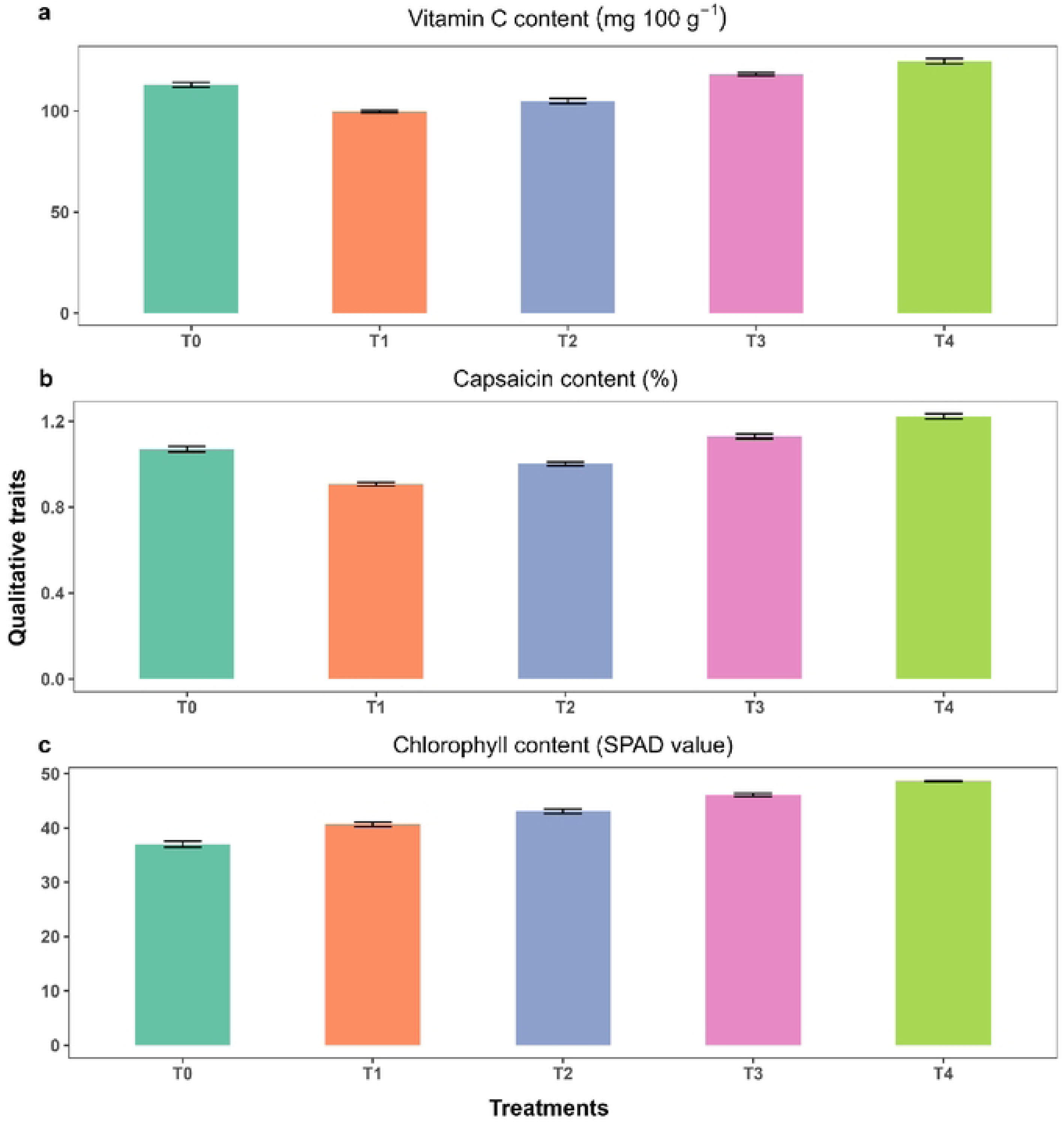
Vitamin C, capsaicin content and SPAD chlorophyll index of *Capsicum frutescence* under different fertilization treatments.

### 3.6 Significant associations between ANPP and vegetative-reproductive traits

ANPP exhibited strong and statistically significant positive relations with both leaf number (*R²* = 0.51, *p* = 0.02) (Fig 10a) and flower number (*R²* = 0.41, *p* = 0.014) (Fig 10b) highlighting that greater vegetative and reproductive growth directly enhanced aboveground biomass accumulation. Among the treatments, only T₄ (organic + ½ RFD) achieved the highest ANPP in relation to its reproductive traits (leaves: *R²* = 0.57, *p* = 0.006 and flowering: *R²* = 0.68, *p* < 0.001).

**Fig 10.**
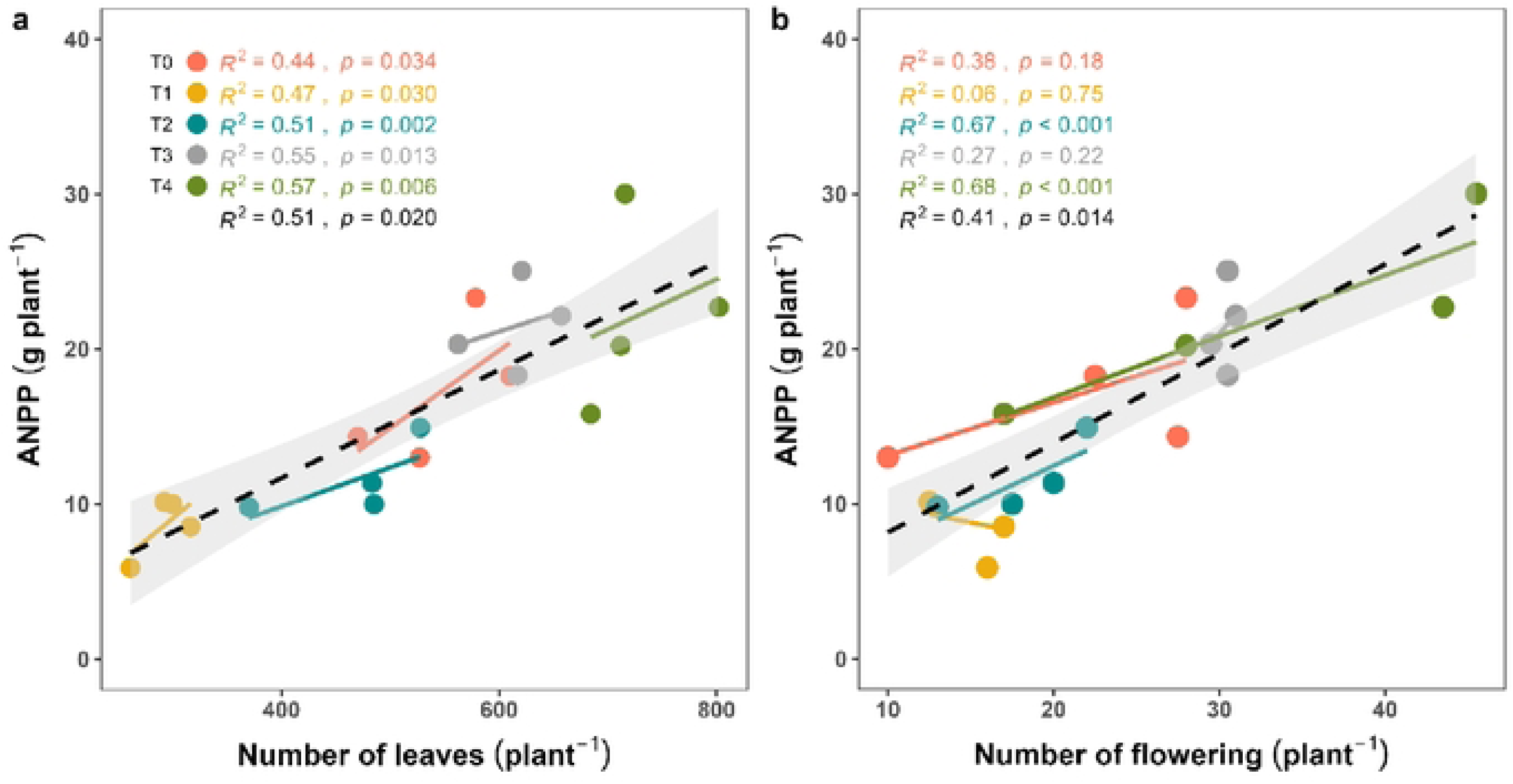
Regression analysis between ANPP and vegetative (leaf number) and reproductive (flower number) traits of *Capsicum frutescence*.

### 3.7 Yield strongly influenced by vegetative and reproductive structures

A positive linear relationship was observed among yield, number of leaves (*R²* = 0.34, *p* = 0.002) (Fig 11a) and the number of flowering plant^-1^ (*R²* = 0.42, *p* = 0.037) (Fig 11b), indicating their strong predictive value for yield performance. Among treatments, T₂ showed the significantly higher relation between leaf number versus yield (*R²* = 0.80, *p* < 0.001), as well as flowering versus yield (*R²* = 0.82, *p* < 0.001) respectively. T₁ also showed a moderate but significant relationship between yield and reproductive traits (leaves: *R²* = 0.40, *p* = 0.041 and flowering: *R²* = 0.33, *p* = 0.046). In contrast, the yield of T₄ did not show a significant correlation with plant reproductive traits (leaves: *R²* = 0.12, *p* = 0.65 and flowering: *R²* = 0.28, *p* = 0.41).

**Fig 11.**
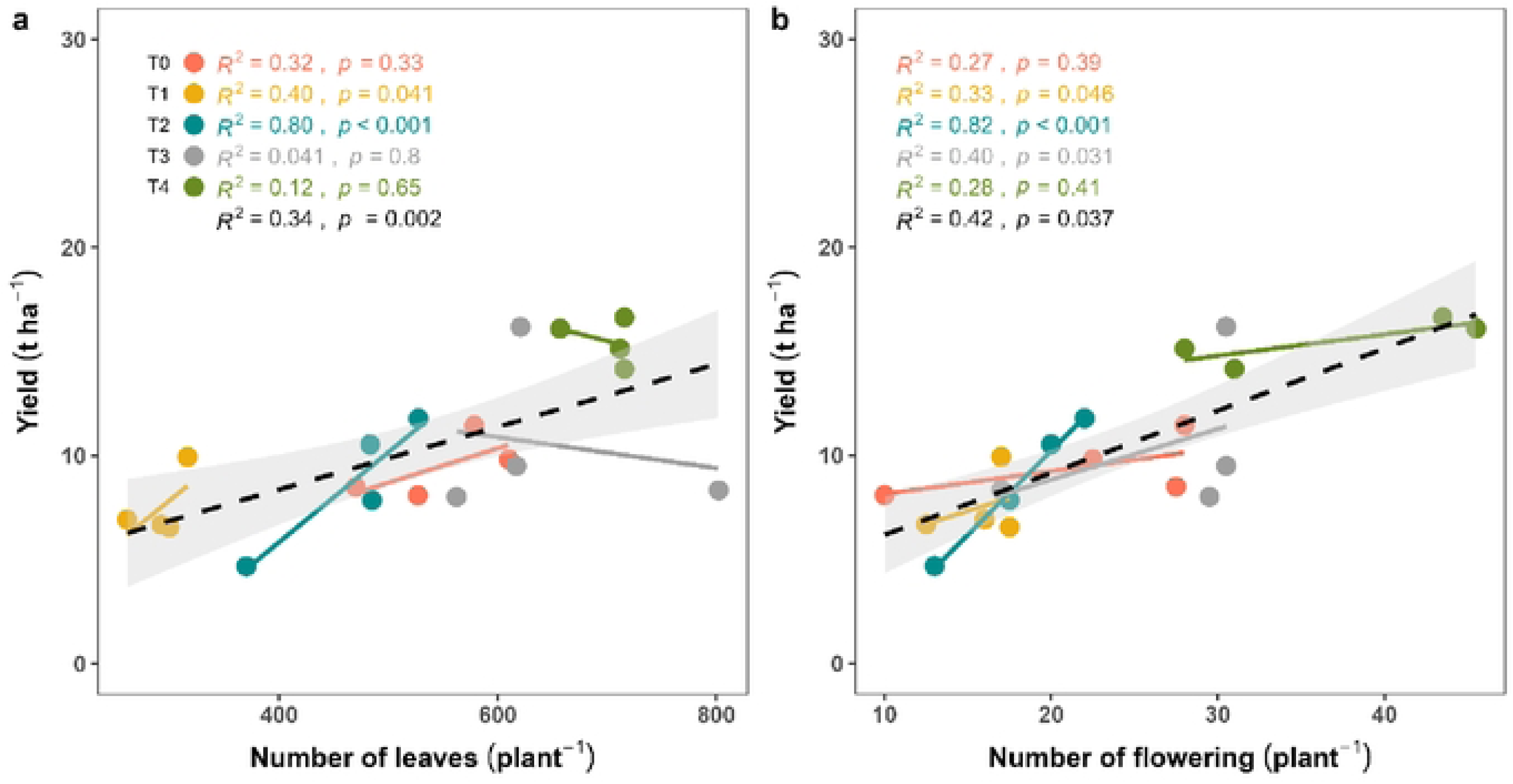
Regression analysis showing relationship between fruit yield and vegetative-reproductive traits (leaf and flower number) in *Capsicum frutescence*.

## 4. Discussion

### 4.1 Effect of mixed and mono fertilization on yield-contributing traits of chili

The vegetative traits-specifically plant height and leaf number-showed significant differences among the fertilization treatments, with mixed fertilization (T₄) consistently outperforming mono strategies at all observation stages. These traits are important yield-contributing characteristics because they influence photosynthetic efficiency, carbon assimilation and plant vigor [26,55]. The superior vegetative performance under T₄ can be attributed to the dual advantage offered by mixed fertilization, firstly, immediate nutrient availability through inorganic fertilizers and secondly, gradual nutrient release through organic amendments like cow dung and *Leucaena* leaf litter. These observations are in direct alignment with the findings presented in plant height (Fig 3) and leaf number (Fig 4), where T₄ displayed markedly higher values at all stages. Previous studies have corroborated the beneficial effects of integrated nutrient management (INM) in promoting structural growth. For instance, Malik et al. [23] and Zahid et al. [22] reported improved plant height, branching and canopy development in sweet pepper and cucumber respectively when organic manure (e.g. poultry manure) was combined with reduced synthetic inputs. These findings are consistent with earlier studies indicating that the synergistic application of organic and inorganic fertilizers enhances vegetative vigor due to improved nutrient availability, microbial activity and soil structure [15,16]. The reproductive traits-number of flowers and fruits per plant were also significantly enhanced under mixed fertilization (Figs 5 and 6). In our experiment, at 60 DAT T₄ plants exhibited nearly double the number of fruits compared to the control and mono- organic treatments (Fig 6c). The increased flower formation and fruit set under integrated fertilization suggest a positive hormonal and nutritional environment conducive to reproductive development. According to Sharma & Mittra, [33], the presence of both macro and micronutrients in balanced proportions facilitates flowering and fruiting by supporting enzyme activity and hormonal synthesis. Although we did not measure the mechanism behind the findings, however, considering other experiment, we assumed that the presence of organic matter such as *Leucaena* litter enhances rhizospheric activity and microbial interactions that release essential growth hormones like cytokinins and auxins, further promoting flower retention and fruit development [18,19]. Such synergistic outcomes cannot be achieved through mono fertilization, especially when nutrient availability is either too rapid (in the case of synthetics) or too slow (in the case of organics). Hence, mixed fertilization offers a more buffered and physiologically aligned nutrient release pattern to support yield-contributing growth traits.

### 4.2 Effect of mixed and mono fertilization on NPP and fruit quality

The net primary productivity (NPP), measured as the sum of aboveground (ANPP) and belowground (BNPP) biomass, was significantly greater under mixed fertilization strategies than mono treatments. T₄ recorded the highest total NPP indicating that integrated fertilization not only enhances shoot growth but also supports robust root development (Fig 7c). These findings are consistent with earlier work by Jiang et al. [20] and Haque et al. [56], who observed increased biomass accumulation under integrated nutrient management (INM) treatments due to improved nutrient use efficiency and soil nutrients dynamics. In our findings, increased BNPP under mixed fertilization could be linked to enhanced root proliferation, root rhizosphere, microbial decomposition and nutrient allocation particularly carbon and nitrogen which favored positively to root biomass [57]. Enhanced BNPP under mixed treatments suggests a healthy root system that can efficiently explore the soil matrix for water and nutrients. Similar scenario was observed in our experiment where mixed fertilization (e.g. *Leucaena* leaf litter & reduced synthetic fertilizer) response more positively (3.93 g plant^-1^) in case of BNPP of chili compared to control (3.25 g plant^-1^) and inorganic (1.95 g plant^-1^) fertilizers (Fig 7b). Root development is strongly influenced by the physical and biological properties of the soil, which are significantly improved by the addition of organic matter [40,58]. Organic inputs improve soil texture, moisture retention and microbial colonization-all of which contribute to enhanced root growth [59]. Moreover, deeper and denser root systems not only increase BNPP but also contribute to long-term soil organic carbon storage and soil health (REF) [60]. Fruit yield was significantly enhanced under mixed fertilization with T₄ achieving the highest yield (15.50 t ha⁻¹), which is 63.9% increase over the control (Fig 8). This improvement is attributed to the combined benefits of rapid nutrient availability from inorganic fertilizers and sustained nutrient release from organic inputs, which together optimize nutrient use efficiency and support continuous reproductive development [16,23]. The superior yield of chili under T₄ also aligns with findings by Jiang et al. [20] and Gokul et al. [26], who reported enhanced biomass and fruit productivity under integrated nutrient management (INM). Organic amendments further improve soil structure and microbial activity, promoting root development and nutrient uptake [19,56]. These results confirm that mixed fertilization strategy boosts up chili production. In terms of fruit quality, the T₄ treatment again showed outer performance. Vitamin C concentration, capsaicin content and SPAD chlorophyll index were highest under mixed fertilization (Fig 9). Vitamin C and capsaicin are key quality indicators in chili, with implications for both nutrition and market value. Higher levels of these biochemical compounds under T₄ suggest that integrated fertilization supports optimal metabolic functioning (MF) of the plant. Although we did not measure the effect of metabolic functioning of chili but considering other results, we assume that mixed fertilizer may enhances the metabolic function in the plants. The increased vitamin C under mixed fertilization is likely due to improved micronutrient availability, particularly potassium and magnesium which play a critical role in ascorbate biosynthesis [24]. Similarly, capsaicin synthesis is known to be enhanced under balanced nitrogen regimes, which can be achieved through mixed fertilization [23]. The enhanced SPAD chlorophyll index indicates improved nitrogen assimilation and photosynthetic activity, further corroborating the physiological advantages of mixed input regimes [26]. Overall, the results confirm that mixed fertilization not only boosts biomass production (Fig 7) but also enhances fruit quality (Fig 9) offering a dual benefit for sustainable chili cultivation.

### 4.3 Relationships among morphological traits, biomass accumulation and fruit yield

Regression analysis revealed significant positive relationships between vegetative and reproductive traits and both ANPP and fruit yield. Specifically, ANPP was positively related with leaf number (*R*² = 0.51, *p* = 0.020) (Fig 10a) and flower number (*R²* = 0.41, *p* = 0.014) (Fig 10b) respectively. Similarly, fruit yield was positively linked-with both vegetative (leaf count) and reproductive (flower count) traits (Fig 11). Monteith [29] proposed that crop yield is a function of intercepted radiation, photosynthetic efficiency and biomass partitioning-all of which are mediated by structural traits like leaf area and canopy architecture. Furthermore, Tilman et al. [12] and Agele et al. [30] have emphasized the role of nutrient balance in optimizing the source-sink relationship in crops, thereby improving yield. Importantly, this study demonstrated that even partial replacement of synthetic fertilizers with organic sources as in T₃ (leaf litter 93 g pot^-1^ + ½ RFD) and T₄ (leaf litter 186 g pot^-1^ + ½ RFD) can significantly improve biomass and yield without compromising physiological efficiency (Figs 7, 8, 10 and 11). This has major implications for fertilizer management in resource-constrained settings where synthetic inputs are expensive or environmentally detrimental [61]. The strong positive correlations observed in this study indicate that mixed fertilization can serve as a strategy for improving nutrient use efficiency (NUE) and physiological productivity [62]. The integrative approach to nutrient supply-combining the strengths of fast-acting inorganic fertilizers and the soil-enhancing qualities of organics-creates a robust production environment where plants can achieve their full physiological and genetic potential. This not only leads to higher yields but also promotes long-term sustainability and resilience of the farming system (REF) [63].

## 5. Conclusions

The findings of this study demonstrate that mixed fertilization strategies exert a more substantial positive influence on chili (*Capsicum frutescens*) growth, productivity and quality than mono fertilization approaches. Plants grown under integrated nutrient management, combining organic inputs like *Leucaena leucocephala* leaf litter with reduced levels of inorganic fertilizers uses, showed markedly better vegetative development, reproductive output and physiological vigor compared to those under sole organic or inorganic treatments. Three key outcomes emerged from this research. **Firstly,** vegetative traits such as plant height and leaf number were consistently enhanced under mixed fertilization, suggesting improved nutrient availability and plant health. **Secondly,** aboveground and belowground biomass accumulation (represent the ANPP and BNPP) were notably higher in integrated treatments, indicating superior carbon allocation and net primary productivity. **Finally,** essential fruit quality attributes including vitamin content, capsaicin concentration and chlorophyll index-were significantly improved under mixed fertilization, highlighting its positivity to metabolic and biochemical functions.

Overall, the study underscores that mono fertilization-whether organic or inorganic-offers limited benefits when compared to combined mixed-mono approach. The synergistic effect of mixed fertilization not only enhances plant performance and yield but also supports soil fertility and sustainability. Hence, integrated nutrient management should be promoted as a climate-resilient and resource-efficient practice for chili cultivation in Bangladesh and other geographic areas with similar agroecological environment. Future studies should focus on long-term field trials along with others organic amendments to assess soil health, yield stability and at the same time the environmental impacts of such fertilization practices across diverse agroecological conditions.

## Acknowledgments

This research was funded by the Sher-e-Bangla Agricultural University Research System (SAURES) grant no. SAU/SAURES/2022/110(20) and Ministry of Science and Technology, Bangladesh, through a special research allocation grant (Grant No: SRG-221311(BS)/2022-23).

## Author contribution

Conceptualization: Hossain Md Dalim, Md. Golam Jilani Helal

Data curation: Dr. Md. Shahariar Jaman, Hossain Md Dalim, Minhazul Kashem Chowdhury Formal analysis: Hossain Md Dalim, Dr. Md. Shahariar Jaman

Funding acquisition: Md. Golam Jilani Helal, Dr. Md. Shahariar Jaman Investigation: Dr. Md. Shahariar Jaman, Md. Golam Jilani Helal

Methodology: Hossain Md Dalim, Dr. Md. Shahariar Jaman, Md. Golam Jilani Helal, Project administration: Dr. Md. Shahariar Jaman, Md. Golam Jilani Helal

Resources: Md. Mohi Uddin Sujan Chowdury, Shoaib Rahman

Software: Hossain Md Dalim, Md. Golam Jilani Helal, Dr. Md. Shahariar Jaman Supervision: Dr. Md. Shahariar Jaman

Validation: Shoaib Rahman, Md. Ismail Hossain, Sohel Rana Mazumder, Minhazul Kashem Chowdhury

Visualization: Md. Ismail Hossain, Sohel Rana Mazumder, Md. Mohi Uddin Sujan Chowdury Writing – original draft: Hossain Md Dalim

Writing – review & editing: Hossain Md Dalim, Md. Golam Jilani Helal, Dr. Md. Shahariar Jaman, Minhazul Kashem Chowdhury.

